# Analysis of duplication and possible sub-functionalization of wing gene network components in pea aphids

**DOI:** 10.1101/2025.04.30.651519

**Authors:** Omid Saleh Ziabari, Kevin D. Deem, Qingyi Zhong, Jennifer A. Brisson

**Affiliations:** Department of Biology, University of Rochester, Rochester, NY, 14627; Department of Biological Sciences, University of Pittsburgh, Rochester, PA 15260

**Author notes:** These authors contributed equally to this work (Omid Saleh Ziabari, Kevin D. Deem).

**Keywords:** wing, gene regulatory network, dimorphism, pea aphids

## Abstract

A fundamental focus of evolutionary-developmental biology is uncovering the genetic mechanisms responsible for the gain and loss of characters. One approach to this question is to investigate changes in the coordinated expression of a group of genes important for the development of a character of interest (a gene regulatory network). Here we consider the possibility that modifications to the wing gene regulatory network (wGRN), as defined by work primarily done in *Drosophila melanogaster*, were involved in the evolution of wing dimorphisms of the pea aphid (*Acyrthosiphon pisum*). We hypothesize that this may have occurred via changes in expression levels or duplication followed by sub-functionalization of wGRN components. To test this, we annotated members of the wGRN in the pea aphid genome and assessed their expression levels in first and third nymphal instars of winged and wingless morphs of males and asexual females. We find that only two of the 32 assessed genes exhibit morph-biased expression. We also find that three wing genes (*apterous* (*ap*), *warts* (*wts*), and *decapentaplegic* (*dpp*)) have undergone gene duplication. In each case, the resulting paralogs show signs of functional divergence, exhibiting either sex-, morph-, or stage-specific expression. Two gene duplicates, *wts2* and *dpp3*, are of particular interest with respect to wing dimorphism, as they exhibit a wingless male-specific isoform and wingless male-biased expression, respectively. These results supplement our understanding of trends in developmental gene network evolution, such as side-stepping pleiotropic constraint via duplication and sub-functionalization, underlying the emergence of novel phenotypes.

## Introduction

Phenotypic novelties have long fascinated scientists, often involving mysterious and rapid emergence or disappearance of entire structures with long-lasting evolutionary impacts. For example, the novel insect wing and advent of powered flight are frequently cited as a major contributor to the success of the insects (Grimaldi and Engel, 2005). Perhaps counterintuitively, nearly every extant winged insect clade also contains members which have secondarily evolved flightless or even wingless members (Wagner and Liebherr, 1992). Many evo-devo studies over the past several decades have begun to shed light on the molecular mechanisms underlying the origin of the novel wing (Clark-Hachtel and Tomoyasu, 2016; Tomoyasu, 2021; Treidel et al., 2024). However, our understanding of the evolution of secondary wing loss lags significantly behind (Treidel et al., 2024).

Clues to how drastic changes in body structure might evolve at the molecular level emerged from developmental studies in *Drosophila melanogaster* in the 1990’s. In some cases, the ectopic expression of a particular transcription factor, such as *eyeless* (*ey*) or *vestigial* (*vg*), was sufficient to induce the formation of fully formed eyes or wings, respectively, in other parts of the body (see Halder et al., 1995 in regards to *ey*, see Baena-López and García-Bellido, 2003; Maves and Schubiger, 1998; Paumard-Rigal et al., 1998 in regards to *vg*). Likewise, loss of function for these types of “master regulator” transcription factors would lead to complete loss of their associated structure (Quiring et al., 1994 for *ey*, Williams et al., 1991 for *vg*). Understanding how body patterning signals activated master regulators within their proper location to establish field primordia, as well as how downstream growth and differentiation genes were subsequently activated, became of primary importance in developmental biology. In the proceeding decades, massive interest was paid to unveiling the networks of regulatory logic governing the expression of such genes, termed Gene Regulatory Networks (GRNs).

One potentially fruitful avenue for identifying the mechanisms underlying the rapid loss or gain of whole structures within or between species is by studying the structure and deployment of GRNs. In particular, the developmental GRN underlying wing formation in *Drosophila* (wGRN) is one of the best-understood to date (see Tripathi and Irvine, 2022 for a review of the wGRN, see Linz et al., 2023 for a diagram of the network). This has laid the groundwork for analyzing orthologs of these wGRN components in wingless or wing dimorphic species to study the evolution of wing loss

Wing dimorphisms of pea aphids (*Acyrthosiphon pisum*) are an excellent model for studying the evolution of winglessness. Pea aphids produce both winged and fully wingless morphs in two ways: via environmental induction in asexual females (wing plasticity), and under the control of distinct genetic alleles in males (Brisson, 2010). While most aphid species exhibit the asexual female wing plasticity, male wing dimorphisms are rare and are likely a transition state between monomorphic winged or wingless male states for a species (Saleh Ziabari et al., 2023). Thus, pea aphids allow us to study not only wing loss in the context of plasticity, but also during the process in which winglessness becomes genetically controlled and eventually fixed in a clade. Recent advances have been made in our understanding of the molecular mechanisms governing wing dimorphisms in aphids (Deem et al., 2024; Li et al., 2020; Saleh Ziabari et al., 2025). Still, much remains to be learned regarding the modulation of GRNs necessary to achieve the wingless phenotype, or if members of the wGRN identified in *Drosophila* are involved.

The presence of winged aphid morphs in asexual females of nearly all aphid species requires that some version of the wGRN is intact and functional, and thus, loss of wGRN components is an unlikely culprit for the evolution of winglessness in this clade. Further, most wGRN components are involved in numerous other developmental processes, so there is little expectation that they would be completely repressed in a wingless individual. As a result, these types of pleiotropic genes may evolve novel expression domains by gaining new, context-specific regulation which does not perturb gene function in other contexts (Kryuchkova-Mostacci and Robinson-Rechavi, 2016). However, this process is expected to be somewhat slow to evolve relative to other mechanisms, and is difficult to study, usually requiring context-specific transcriptomic, epigenetic, and enhancer-activity information (Deem and Brisson, 2024; Kryuchkova-Mostacci and Robinson-Rechavi, 2016).

There are alternative evolutionary trajectories that may be assessed by comparative analysis of more broadly available data, such as whole-body transcriptomes. Although expected to be unlikely, we have yet to rule out the possibility that systemic repression of one or more pleiotropic wing genes produces all or many of the characters associated with the wingless aphid morphs. As mentioned above, the expected rarity of such an event stems from the requirement that negative pleiotropic effects must be outweighed by the fitness advantage of the novel characters. Gene duplication followed by sub- and neo-functionalization presents another trajectory which shows more promise, due to bypassing the barrier of negative pleiotropic effects (Kryuchkova-Mostacci and Robinson-Rechavi, 2016; Kuzmin et al., 2022; Ohno, 1970).

Within the context of pea aphid wing dimorphisms, both of these possibilities can be assessed using existing whole-body RNA-Seq data from winged and wingless male and female morphs.

In both cases, the single-copy pleiotropic wing genes, as well as those that have duplicated and potentially become morph-specific, should exhibit markedly different expression between morphs. We re-analyzed RNA-Seq data (Saleh Ziabari et al., 2025) taken from male and female winged and wingless morphs in the first and third nymphal instars before and after degeneration of the wing primordia (Ogawa and Miura, 2013). We identified differentially expressed genes within the wGRN and manually curated duplicates. These data were leveraged to evaluate what roles the systemic repression of pleiotropic wGRN single-copy orthologs, or sub-functionalization of duplicates, may have played in the evolution of winglessness in aphids.

## Materials and Methods

### Ortholog Identification

To identify pea aphid orthologs of *D. melanogaster* wing development genes, we performed a BLASTP (Camacho et al., 2009) search using the *D. melanogaster* protein sequence querying *A. pisum* (taxid: 7029) in the RefSeq protein database (O’Leary et al., 2016). Hits with an E-value less than 1e-25 were used in a BLASTP search querying *D. melanogaster* (taxid: 7227) RefSeq proteins. We assigned 1:1 orthology to sequences where the BLASTP and the reciprocal BLASTP results matched. For the sequences that did not have matching results or had multiple hits (*apterous, warts*, and *decapententaplegic*), we performed multiple alignments with structurally related protein sequences from *D. melanogaster, A. melifera, T. castaneum*, and pea aphids, and built phylogenetic trees to confirm orthology relationships. To build the trees, we first used MAFFT (Katoh and Standley, 2013) to align the sequences, then built trees using IQ-TREE (Minh et al., 2020) with 1000 bootstrap replicates.

### RNA-Seq Analyses and Expression Analysis

We collected first and third instar nymphs of winged and wingless asexual females and males. A complete description of the samples can be found in Saleh Ziabari et al., 2025. The read accession numbers for RNA-Seq samples used in this study are SRR32079974, SRR32079975, SRR32079976, SRR32079978, SRR32079989, SRR32080000, SRR32080006, SRR32080007, SRR32080011, SRR32080012, SRR32080013, SRR32080014, SRR32080015, SRR32080016, SRR32080017, SRR32080021, SRR32080022, SRR32080023, SRR32080024, SRR32080025, SRR32080026, SRR32080027, SRR32080028, SRR32080029, SRR32080030, SRR32080031, SRR32080032, SRR32080033, SRR32080034. We used the same RNA-Seq data, filtering criteria, analysis, and mapping approaches described in (Saleh Ziabari et al., 2025). Briefly, this involved removing low quality reads, mapping reads with STAR (2.7.3a) (Dobin et al., 2013) to the pea aphid v3 reference genome (Li et al., 2019) with standard parameters, except increased mapping stringency (as described in Saleh Ziabari et al., 2025). We used featureCounts (2.0.3) within the Rsubread package (Liao et al., 2019) with default parameters to get raw count data, and DESeq2 (Love et al., 2014) to normalize and analyze differential gene expression data (see **Table S1** for normalized read counts at each wing gene for each sample). We used a multi-factor design that incorporated “count ∼ sex + stage + wing” and specified the contrast on winged versus wingless comparisons. We separated female and male analyses as recommended by the DESeq2 vignette due to scale of differences in PCA clustering, except for when looking for wing regulatory gene expression differences: we analyzed the full and partial models. We also excluded one male winged third instar sample by outlier criteria of PCA clustering and count frequency distribution data. We used the R package ComplexHeatmap (Gu et al., 2016) to draw a heat map of the expression data with genes organized by pathways and samples organized by sex, instar, and wing morph. Finally, two methods were used to detect novel splice junction sites: 1) StringTie (Pertea et al., 2015) assembly of isoforms from RNA-Seq data (see **Supplemental data files** for genomic coordinates), and 2) the two-step mapping STAR pipeline, as in Saleh Ziabari et al., 2025. Splice junctions were considered supported if found in all replicates.

## Results

### Pea aphid wing network genes

We focused on 32 wGRN genes as described in *Drosophila melanogaster* (**Table 1**). These include patterning genes upstream of the wGRN, the wing master gene *vestigial* (*vg*) and its ancient paralog *Tondu-domain-containing Growth Inhibitor* (*Tgi*, to help identify the *bona-fide vg*), additional downstream genes within the wGRN, and the Hox genes expressed within the thorax. Many of these genes are important for the activation, spread, and maintenance of *vg* expression in the nascent wing. Expression of *vg* is initiated within the wing primordia downstream of the dorsoventral patterning axis involving *apterous* (*ap*), *fringe* (*fng*), *Delta* (*Dl*), *Serrate* (*Ser*), *Notch* (*N*), and *wingless* (*wg*) (de Celis et al., 1996; Kim et al., 1996; O’Keefe and Thomas, 2001; Williams et al., 1994). Maintenance and induction of *vg* throughout the rest of the growing wing occurs though four mechanisms: **1)** anteroposterior patterning involving *engrailed* (*en*), *patched* (*ptc*), *hedgehog* (*hh*), *cubitus interruptus* (*ci*), *decapentaplegic* (*dpp*), *thickveins* (*tkv*), and *Mothers against decapentaplegic* (*Mad*) (Kim et al., 1996; Zecca et al., 1995; Zecca and Struhl, 2021), **2)** induction of *vg* in pre-wing cells recruited from its periphery in a feed-forward mechanism involving *fat* (*ft*), *dachsous* (*ds*), *dachs* (*d*), *warts* (*wts*), *yorkie* (*yki*), and *scalloped* (*sd*), **3)** continuous *wg* signaling from the dorsoventral boundary and the periphery of the developing wing, and **4)** autoregulation of *vg* by the Vg::Sd transcription factor complex (Zecca and Struhl, 2021, 2010, 2007). Other wing patterning genes include *optomoter blind* (*omb*), *spalt-major* (*salm*), *ventral veins lacking* (*vvl*), and *brinker* (*brk*) (anteroposterior patterning), as well as *distalless* (*dll*), *homothorax* (*hth*), *nubbin* (*nub*), *tiptop* (*tio*), and *teashirt* (*tsh*) (proximodistal patterning) (Tripathi and Irvine, 2022). In addition, the Hox genes *Sex combs reduced* (*Scr*), *Antennapedia* (*Antp*), and *Ultrabithorax* (*Ubx*), were included for specifying thoracic segmental identity and interacting with the wGRN to produce either a wingless first thoracic segment, forewings, or hindwings, respectively (Fang et al., 2022; Lewis, 1978; Tomoyasu et al., 2005).

**Table 1:**
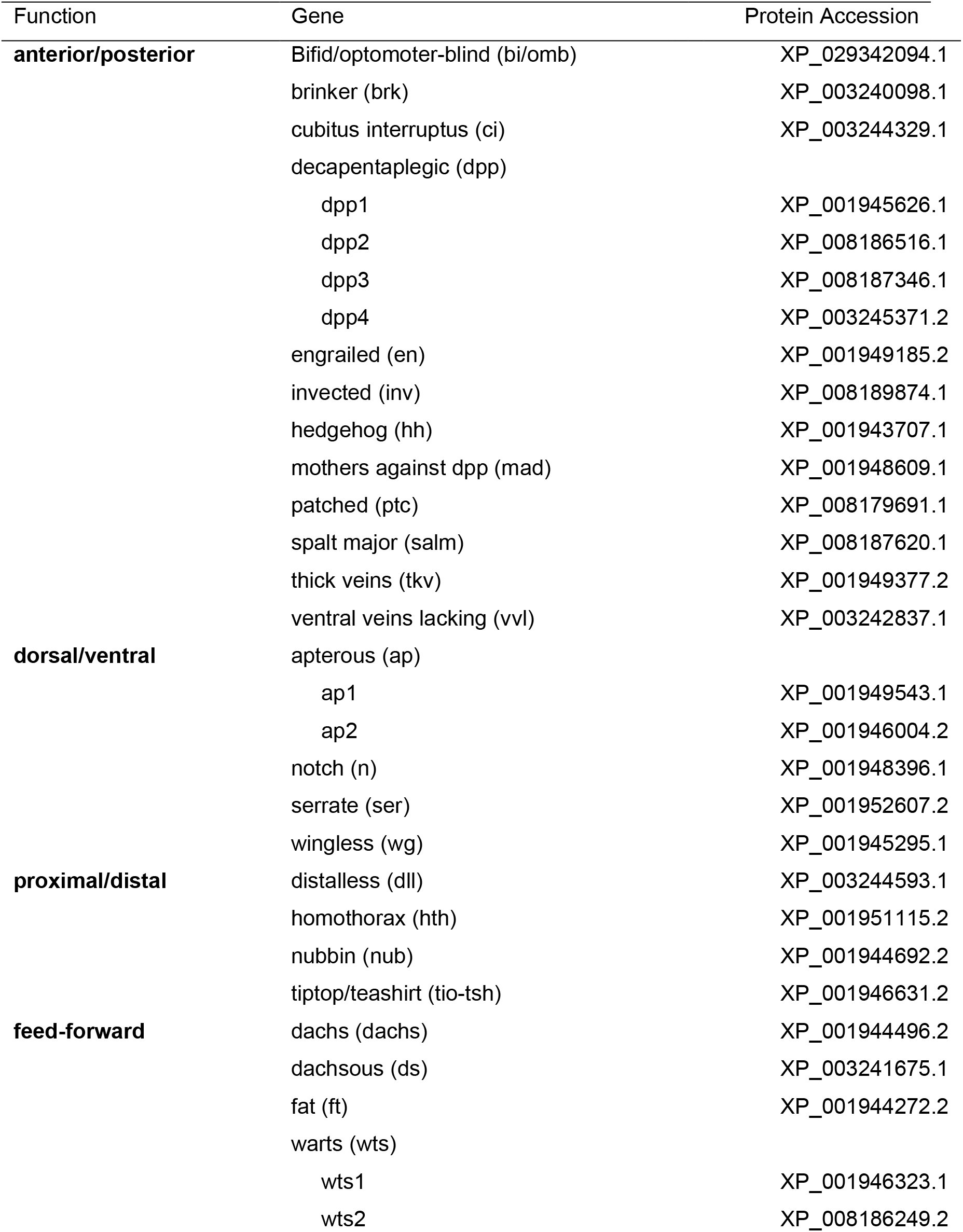
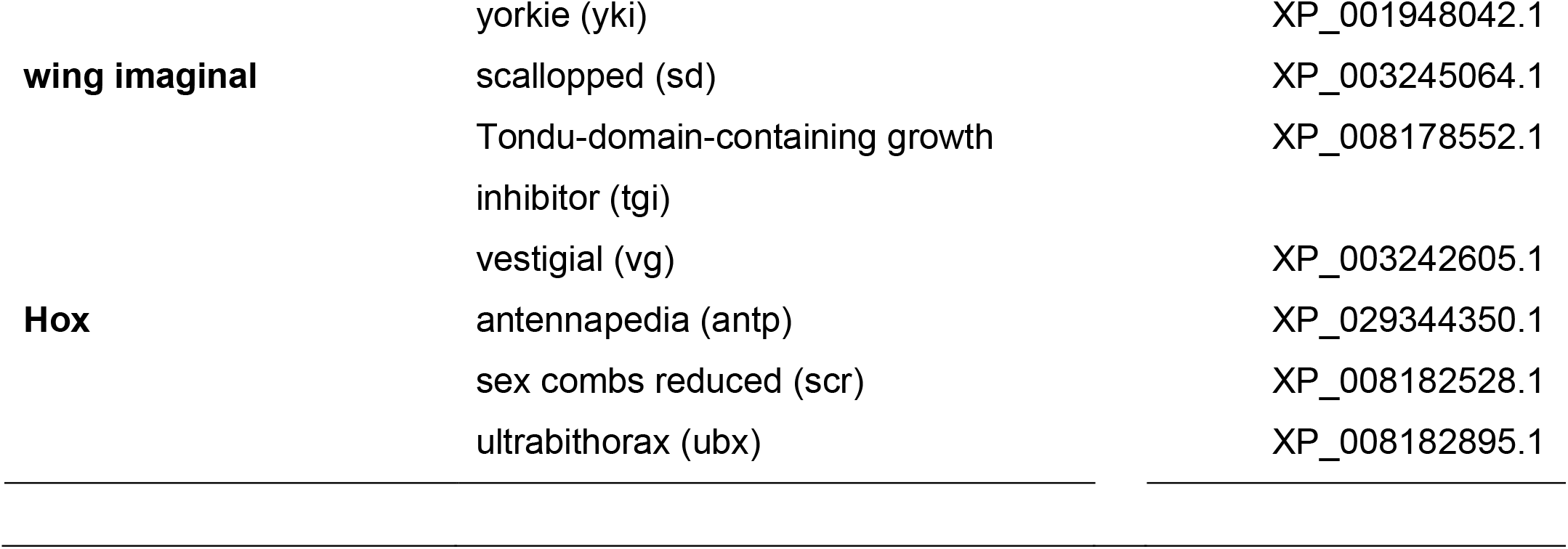
Pea aphid wing gene regulatory network in pea aphids. The designations of patterning functions are bolded in the first column, the gene names and their abbreviations in the second column, and their associated NCBI protein accessions in the final column. Indented genes represent detected duplications.

### Wing gene expression level differences between pea aphid wing morphs

We asked if genes within the wing regulatory network showed broad differences in gene expression levels between pea aphid wing morphs across two nymphal instars (first and third; pea aphids have four nymphal instars prior to adulthood), within both males and females. To do this, we categorized wing regulatory network genes by their function (as in Table 1) and then manually ordered samples by sex and nymphal instar. Despite the inclusion of canonical wing network genes, we observed no overall pattern of differential expression levels between winged and wingless morphs of either sex with these genes (**Fig. 1A**).

**Figure 1:**
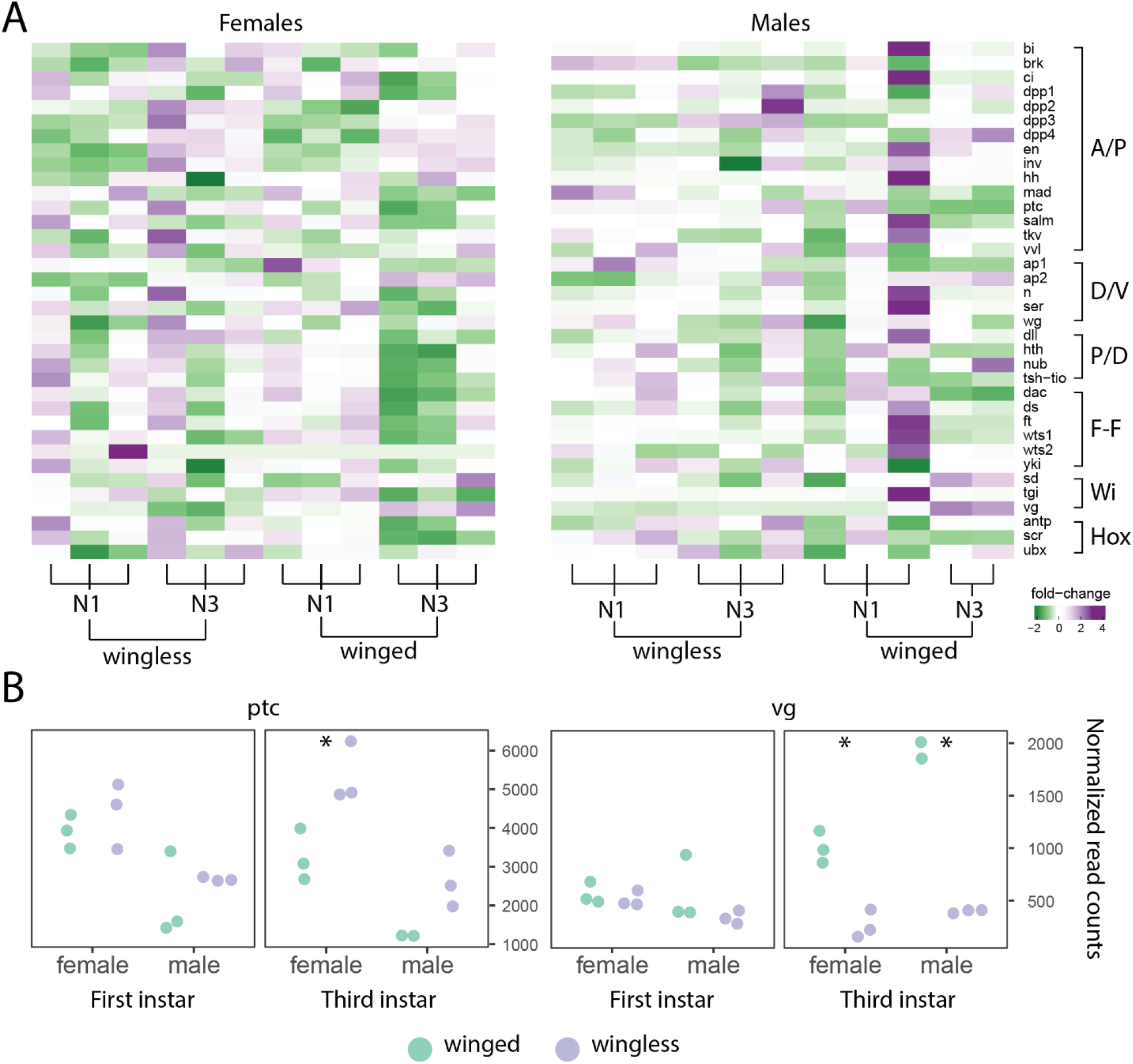
Wing gene expression levels in winged compared to wingless morphs. (A) Heatmap showing the fold-change expression differences in a model contrasting winged and wingless samples in females (left) and males (right) separately. The samples (columns) are ordered with respect to wing morph and instar stage, while the genes (rows) are in the same order as **Table 1**, including designations of patterning functions (A/P: anterior/posterior; D/V: dorsal/ventral; P/D: proximal/distal; F-F: feed-forward; Wi: wing imaginal; and Hox: as Hox). Positive fold-change (purple) corresponds to higher expression levels in wingless morphs while negative fold-change (green) corresponds to higher expression levels winged morphs. The column sample information is given by “WL” and “W”, referring to *wingless* and *winged* samples, while “N1” and “N3” corresponds to nymphal stage one and nymphal stage three, respectively. (B) The normalized read counts from each instar (first and third) and sex (female and male) for the two significantly differentially expressed genes, *patched* (*ptc*) and *vestigial* (*vg*). Each dot corresponds to the sequenced library sample, where morph is shown as green (winged) and purple (wingless). Asterisks denote significance in a secondary statistical test on each morph pair for each sex and instar using a Student’s T-test, *p<0.05.

Environmentally induced winglessness occurs in females and genetically determined winglessness occurs in males, so sex-biased expression may indicate differences in the two mechanisms. We therefore asked if each gene’s expression differed significantly between winged and wingless morphs of either sex. We found that *vg* and *ptc* exhibited expression differences in our full multi-factor model with a winged versus wingless contrast (*vg*: log_2_FC = 1.20, p_adj_ = 0.043, *ptc*: log_2_FC = -0.53, p_adj_ = 0.003; see **Table S2**), while only *vg* was significant when sex was separated in the analysis (log_2_FC=1.04, p_adj_ = 0.047 for females, log_2_FC = 1.46, p_adj_ = 0.004 for males; see **Tables S3-4**) as recommended when the variance between factors is substantial. We then examined differential expression of these genes within each instar, between morphs and sexes, outside of the multi-factor model. Specifically, within the third instar, *ptc* exhibited higher expression in the wingless morph of both sexes, although only significantly higher in females (Student’s 2-tailed T-test, p = 0.025 for females, p = 0.078 for males; **Figure 1B**), whereas *vg* showed significantly higher expression in the winged morphs of both sexes (p = 0.003 for females, p = 0.029 for males; **Fig. 1B**).

### Expression differences among wing gene paralogs

To further examine if gene duplication and sub-functionalization of wGRN components may have played a role in evolution of the wingless phenotype, we searched for paralogs among these 32 wGRN components. We re-annotated these genes to identify new isoforms within each duplicate using StringTie (Pertea et al., 2015) and the nymphal RNA-Seq dataset (Saleh Ziabari et al., 2025) (see **Supplemental data files** for genomic coordinates). Then, for each, we explored potential functional divergence as evidenced by differences in expression levels among stages, morphs, or sexes at the whole gene or isoform level. Of these genes, 29 of the 32 had one-to-one reciprocal BLAST hits between pea aphids and *Drosophila melanogaster*, while three genes [*apterous* (*ap*), *warts* (*wts*), and *decapentaplegic* (*dpp*)] yielded multiple gene copies in pea aphids (**Table 1**). We identified two paralogs of *apterous*, which exist as a tandem duplication 130 kb apart on autosome 1. We found two copies of the gene *wts* in the genome, each on separate autosomes. Four *dpp* paralogs exist on multiple chromosomes, with one pair on the X chromosome separated by 6.6 Mb.

For pea aphid paralogs, we created protein trees using orthologs from *Drosophila melanogaster, Apis mellifera*, and *Tribolium castaneum* to roughly resolve the lineage in which the duplicates arose or were lost. We included other members of their respective protein family for each of these trees. The tree containing the two pea aphid *ap* paralogs and other LIM homeobox transcription factors like *Arrowhead* (*Awh*), *Lim3* (*LIM3*), and *tailup* (*tup*) supports the *ap* duplication being older than the common ancestor of the included taxa, with a single loss of an *ap* duplicate in *D. melanogaster* (**Fig. 2A)**, consistent with a previous report (Brisson et al., 2010). The protein tree for the two *wts* paralogs included the related serine/threonine kinases *tricornered* (*trc*) and *ghengis khan* (*gek*). Close clustering of *Ap-wts1* and *Ap-wts2* indicates that the duplication event occurred after the lineage leading to aphids split form the lineage leading to the holometabola **(Fig. 2B)**. The protein tree with the *dpp* paralogs included both branches of TGF-β extracellular ligands: the BMP branch [(*screw* (*scw*), *glass bottom boat* (*gbb*), *maverick* (*mav*)] and Activin branch [*dawdle* (*daw), Activin-b* (*act-b)*, and *myoglianin (myo*)]. This tree strongly supports that the three additional *dpp* duplicates are specific to the lineage leading to aphids in the same manner as the *wts* paralogs (**Fig. 2C**).

**Figure 2:**
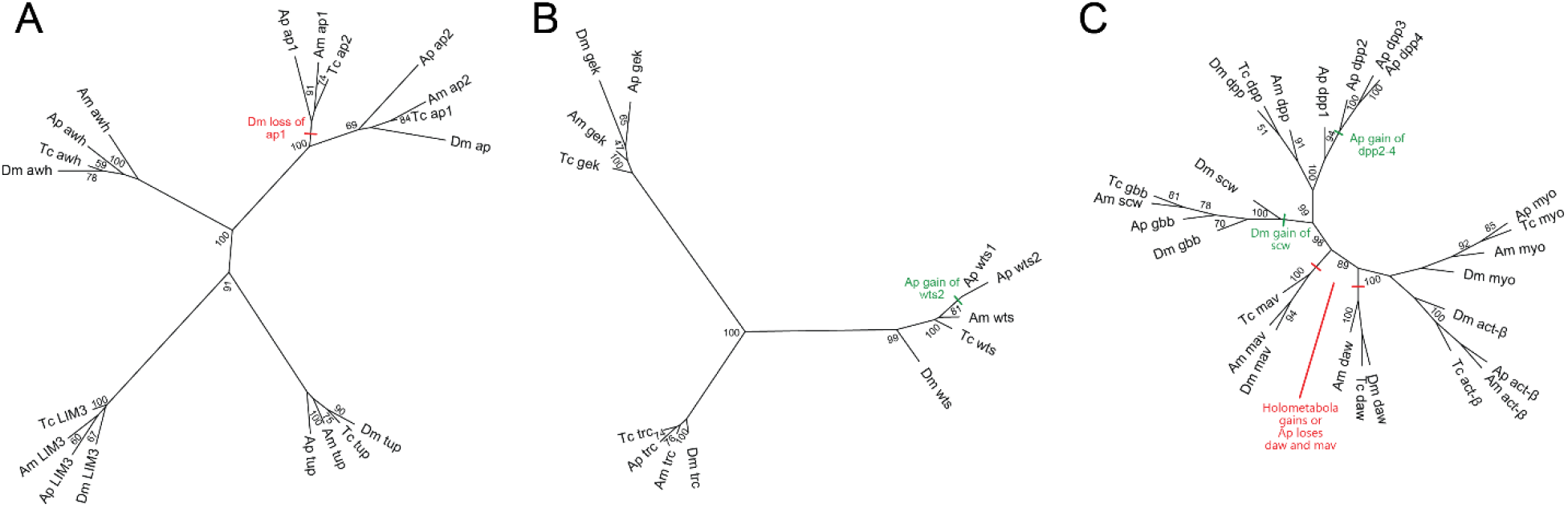
Protein trees of detected paralogs of the wing regulatory network of pea aphids. All protein trees include orthologs from *Acyrthosiphon pisum* (Ap), *Drosophila melanogaster* (Dm), *Apis mellifera* (Am), and *Tribolium castaneum* (Tc). (A) The protein tree for the two *ap* paralogs in pea aphids, including other related LIM homeobox transcription factors: *Arrowhead* (*Awh*), *Lim3* (*LIM3*), and *tailup* (*tup*). Red dash indicates a loss of *Dm-ap1*. Note that the Tc *ap* numerals are backwards from their inferred homology. (B) Protein tree for the two *wts* paralogs, including two related serine/threonine kinases *tricornered* (*trc*) and *ghengis khan* (*gek*). Green dash indicates the gain of *Ap-wts2* (C) The protein tree for the four *dpp* paralogs, including other members of the TGF-β type II extracellular ligands including the BMP [(*screw* (*scw*), *glass bottom boat* (*gbb*), *maverick* (*mav*)] and Activins [*dawdle* (*daw*), *Activin*-β (*act*-β), and *myoglianan* (*myo*)] groups. The green dash in *dpp* cluster indicates Ap gain of *Ap-dpp2, Ap-dpp3* and *Ap-dpp4*. Red dashes along the primary branches of *mav* and *daw* indicate either a gain of these genes in the holometabola, or their loss in Ap. The green dash in the main branch of *scw* and *gbb* indicates the gain of *Dm-scw*. Note, clustering of *Am-scw* with *gbb* orthologs of other species suggests that this gene is mis-annotated in the honey bee. Values by each node show the bootstrap support. Branches are transformed proportionally to outline structure.

We next examined expression levels of the pea aphid paralogs. The gene models for the two *ap* paralogs are illustrated in **Fig. 3A**. Broad differences in gene structure include an extra exon (exon 2) in *ap2* with no detectable sequence homology to exons of *ap1*, and *ap1* having ∼30 kb of intronic sequence less than *ap2*. We found that *ap1* decreased in expression from the first to third instar (p = 0.005, Student’s T-test), while *ap2* increased (p-value < 0.001) (**Fig. 3B**). We also observed that *ap1* was expressed at much lower levels than *ap2*, generally (**Fig. 3B**). We did not, however, observe any statistical differences between morphs or sexes. We did detect a truncated transcript with a new promoter inside intron 4 of *ap2* (**Fig. 3C**), but no morph- or sex-specific usage of that transcript.

**Figure 3:**
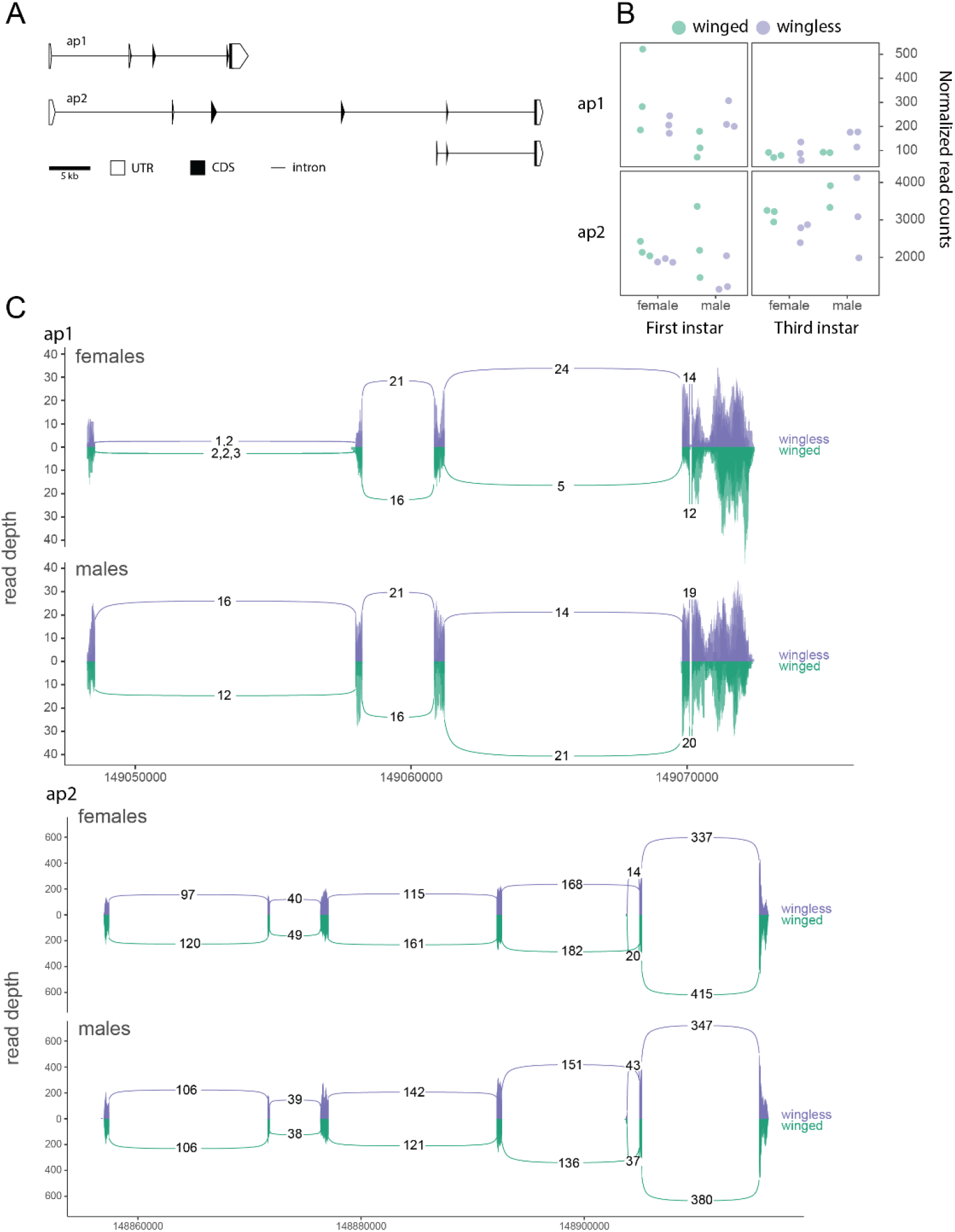
*apterous* (*ap*) paralog structure, expression, and splice variants. (A) Schematic of the gene models for each paralog, and transcript variants detected for *ap1* and *ap2*. Shaded portions illustrate coding region (CDS), while white regions correspond to the untranslated regions (UTR). Connecting lines represent intronic sequence. (B) Gene expression differences between *ap* paralogs from normalized gene expression count data, where *ap1* is shown on the top row and *ap2* is shown in the bottom row. The left column shows first instar nymphal data and the right column shows third instar nymphal data. Females and male samples are separated on the x-axis, and the winged (green) and wingless (purple) morphs are denoted for each replicate. (C) Splice variants are shown where colored bars represent base-pair coverage along the exons, and the arcs connecting each exon represent the splice junction coverage. Numbers represent coverage depth mean across replicates, and are shown when present in all replicates. The top track shows females, and the bottom track shows males, along the genomic interval of *ap1*. Below that, the other splice plot shows the genomic interval of *ap2* (on a different x-axis scale) where the top and bottom still correspond to the female and male samples, respectively. The comma-separated values of the first junction in females shows the low-abundance coverage for each replicate that would otherwise not be shown. Replicates are shown through transparency layers in each plot. Importantly, the winged samples are simply reflected about the y-axis, such that the top portion of each splice track shows the wingless morph expression data (in purple) and the reflection shows the winged samples (in green) on the same scale.

The two paralogs of *wts* also exhibit divergence in their exon-intron structure (**Fig. 4A)**. While neither *wts1* nor *wts2* exhibited any overall morph-biased expression, we detected significantly higher expression of *wts2* in males compared to females in the first (p-value = 0.011) and third (p-value = 0.002) instars (**Fig. 4B**). We also identified one winged and one wingless male-specific isoform for *wts-2*, which are each expressed at a similar level to each other in their respective morph (**Fig. 4C**).

**Figure 4:**
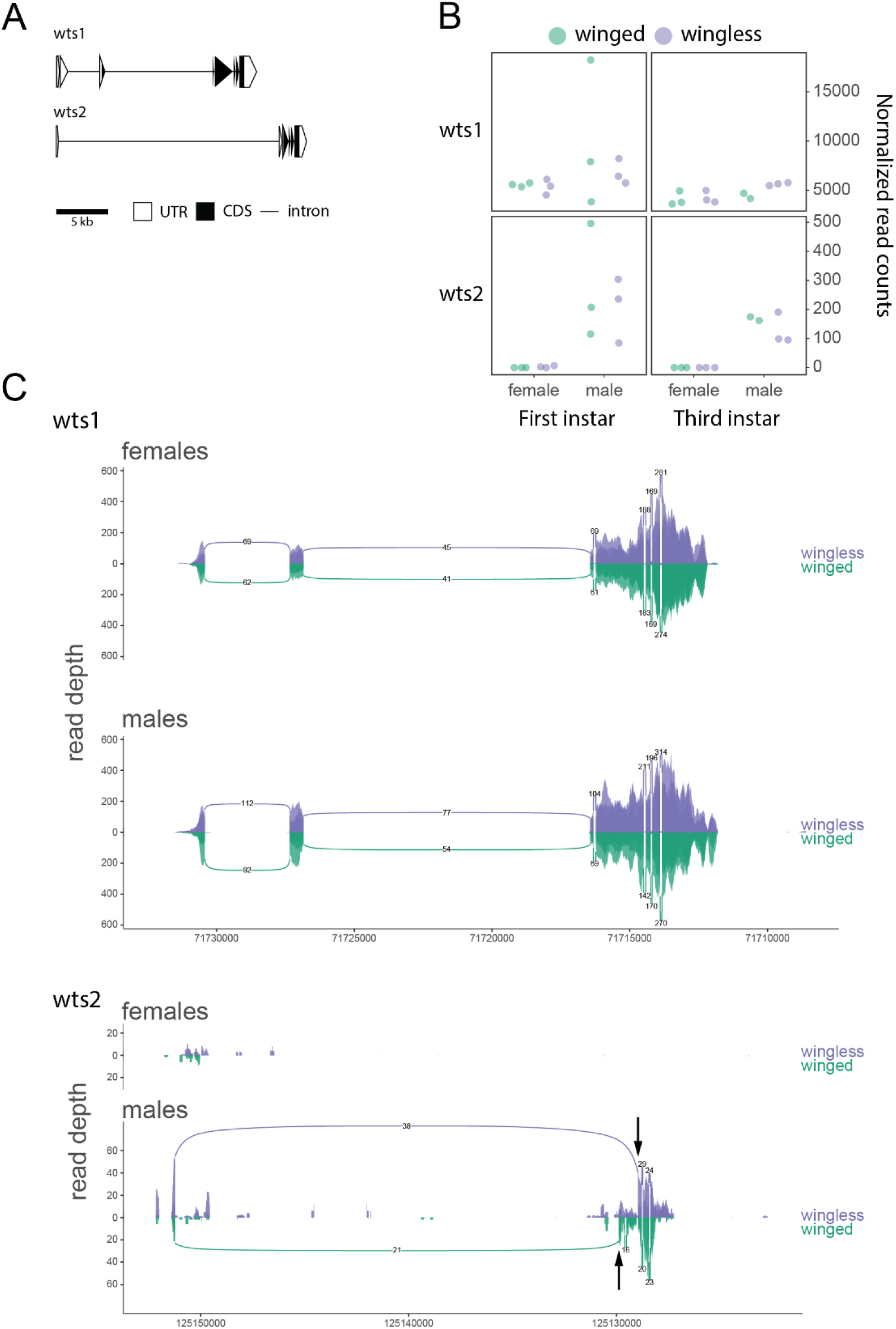
The aphid-specific *wts* paralogs and their expression differences. The gene models, expression profiles, and splicing patterns are shown as in **Figure 3**. (A) The two paralogs of *wts* are shown as gene structure schematics. (B) The normalized read counts across samples for each *wts* paralog. (C) The *wts1* and *wts2* paralogs and their splicing pattern differences are shown. Arrows indicate male morph-specific splice junctions in *wts2*.

The *dpp* paralogs, shown in **Fig. 5A**, exhibited more differences. Although *dpp1*, which had the highest expression levels, exhibited relative stability across stages, sexes, and morphs, the other three paralogs were more dynamic: *dpp2* and *dpp4* were expressed at higher levels in females compared to males at both stages (p < 0.001 for *dpp2*, p = 0.002 for *dpp4*, with first and third instar samples pooled), and *dpp3* had higher levels in wingless relative to winged males (p = 0.011) (**Fig. 5B**). The *dpp* paralogs also presented different splice variants: *dpp3* had a highly expressed, male-specific transcript with two novel exons replacing the first four exons, resulting in a truncated transcript with an intact open reading frame (**Fig. 5C**). We also detected a transcript of *dpp4* that skips exon 2 and retains an open reading frame in another truncated transcript (**Fig. 5C**). Paralog location is associated with sex-specificity, as the two copies on the X chromosome (*dpp3* and *dpp4*) are both male-biased (**Fig. 5C**).

**Figure 5:**
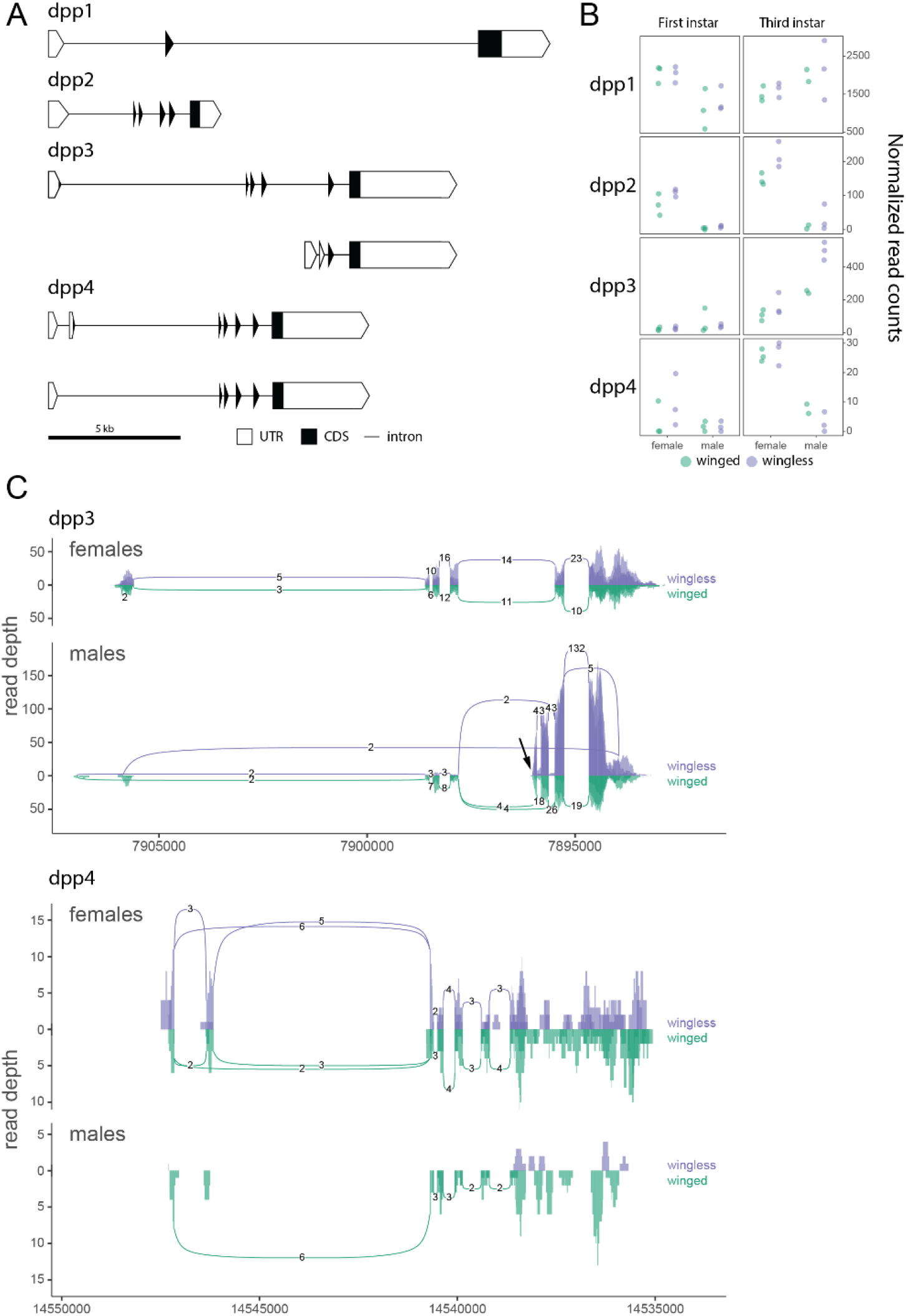
The aphid-specific *dpp* paralogs and their expression differences. The gene models, expression profiles, and splicing patterns are shown as in **Figure 3**. (A) The four paralogs of *dpp* are shown as gene structure schematics and detectable transcript variants. (B) The normalized read counts across samples for each *dpp* paralog in the first and third instar of males and females (C) The splicing differences between *dpp3* and *dpp4*. The transcriptional start site of the male-specific *dpp3* isoform is indicated by a an arrow.

## Discussion

We found that genes in the wing regulatory network are not major contributors gene expression level differences between pea aphid winged and wingless morphs of either sex. Although two genes displayed significantly different gene expression levels between morphs (albeit with low fold changes, log_2_FC = 1.20 for *vg* and -0.53 for *ptc*), the other 30 did not. Thus, it is unlikely that the suite of character differences in wingless morphs is the result of systemic repression of pleiotropic wGRN components. Rather, strong pleiotropic constraint likely precludes evolution of winglessness via substantial loss of wing gene function throughout the body. In the case of *vg*, while the directionality of differential expression aligns with its known function in wing development, it is difficult to ascribe a causal relationship using these data; *i*.*e*., we cannot distinguish between the cases of wings being lost because *vg* expression is lower, versus *vg* expression appearing lower because the wing tissue in which it is broadly and highly expressed is missing.

In the case of *ptc*, determining whether the direction of differential expression aligns with the expectation of wing loss is also quite complicated. The Hedgehog (Hh) signaling pathway is activated when the Hh ligand binds to Ptc to alleviate repression of Smoothened (Smo) (Ingham et al., 2000). Cells that have lost Ptc in the developing *Drosophila* wing tissue lose repression of Smo and exhibit ectopic activation of Hh signaling (Tanimoto et al., 2000; Zhu et al., 2003). This, in turn, represses expression of the Dpp receptor Tkv, and induces loss of p-Mad (the transcriptional effector of Dpp signaling) (Tanimoto et al., 2000). Many important wing genes including *vg, brk, omb, salm*, and *vvl* require proper Dpp signal transduction for their appropriate expression patterns in the nascent wing blade (Tripathi and Irvine, 2022; Zecca and Struhl, 2021). Thus, a decrease in *ptc* expression in winged morphs does not align with known functions in wing development, but only if that loss occurs within regions required to transduce Dpp signaling via the receptor Tkv. Future investigations utilizing histological methods may be able to pinpoint changes in the spatiotemporal expression profiles of *vg* and *ptc* in winged and wingless aphids, helping to clarify potential roles in the differential wing development of these morphs.

Gene duplication and sub-functionalization provides an evolutionary route that bypasses pleiotropic constraints, so we were particularly interested in pea aphid wGRN paralogs. Somewhat surprisingly, we found that only two members, *wts* and *dpp*, duplicated in the lineage leading to the pea aphids. That only two out of the 32 genes examined (6.25%) exhibited aphid lineage-specific duplication is low, given the high levels of gene duplication observed in aphids genome-wide (Li et al., 2019). Paralogs of both *wts* and *dpp* display morph- and sex-biased expression (**Figs. 4, 5)**. The *wts2* paralog exhibited significantly higher expression in males in the first and third instars (**Fig. 4B**), as well as winged and wingless male morph-specific isoforms (**Fig. 4C**). The *dpp* duplicates exhibit signatures of functional differentiation, with sex- and stage-specific expression differences (**Fig. 5B**) as well as expression level differences. Two of the *dpp* paralogs, *dpp3* and *dpp4*, also had different isoforms, with one *dpp3* isoform being highly expressed and male-specific, again suggesting functional divergence.

It is interesting that the two male morph-biased paralogs of *dpp* are the most recently duplicated (**Fig. 2B, Fig. 5B, C**), since duplication of a TGF-β inhibitor *follistatin* has recently been implicated in the evolution of wingless males in this species (Li et al., 2020; Saleh Ziabari et al., 2025). Further, these recently duplicated genes seem to have gained differential regulation in males following translocation to the X chromosome, suggesting this may be a common mechanism for attaining sex-specific regulation. However, being located on the X chromosome is clearly not a requirement for male-biased expression, as evidenced by *wts-2* on autosome 1 (**Fig. 4B, C**). More information regarding the molecular mechanisms of dosage compensation in aphids will be necessary to clarify if and how this sex-biased expression is achieved simply by relocating gene duplicates from autosomes onto the X chromosome.

One confounding factor of our study is that we used whole body sampling, while expectations for wing gene differences would be restricted to wing bud tissue. All pea aphids are born as first instar nymphs with wing buds; it’s only as first and second instars that the wing bud continues to grow in the winged morphs and cell death occurs in the wingless morph (Ogawa et al., 2012; Ogawa and Miura, 2013; Yuan et al., 2023). By the third instar, no remnant of the wing bud remains in wingless morphs (Ogawa et al., 2012; Ogawa and Miura, 2013). Therefore, our gene expression profiling at the first and third instars captured the transcriptomes during these first wing bud differences as well as in a more advanced stages of morph differentiation. However, this method is best suited-for detecting drastic, systemic changes in gene expression throughout the body. As previously mentioned, whole body data fails to provide information regarding minute alterations to context-specific expression profiles (a probable evolutionary trajectory for highly pleiotropic genes; Kuzmin et al., 2022). Thus, we are not able to detect potential context-specific modulation of wGRN components that might occur in the wing primordia of early winged versus wingless nymphs.

This approach was, however, sensitive enough to find wingless-specific promoters of transcript variants in multiple species at *follistatin* (*fs*) paralogs (Saleh Ziabari et al., 2025). The study linked *fs* paralogs to female and male wing dimorphisms in pea aphids, with *fs-2* expression specific to asexual wingless females and *fs-3* expression specific to wingless males. The wingless specificity evolved pre-duplication from alternate promoters in *fs-1*. This is interesting because *follistatin* is an activin inhibitor in *Drosophila*, though it may also inhibit other TGF-β ligands like *dpp* in other species, a focal wGRN member of the current study. A gene like *fs* is outside the typical search criteria of “core” wing network genes, and its involvement illustrates the utility of a more unbiased search for candidate genes.

Our results build upon previous findings (Shigenobu et al., 2010 and Brisson et al., 2010) from when the first pea aphid genome assembly was published (The International Aphid Genomics Consortium, 2010). At that time, the genome was highly fragmented, making it possible that some genes, especially paralogs, were misinterpreted as misassembled scaffolds. Our study confirms the wing gene repertoire of the pea aphid genome. It also adds information about the expression of these genes between morphs and sexes, ultimately showing that *wts* and *dpp* paralogs are functional and likely performing different tasks during development. Finally, this study provides support for the general trends in evolution of 1) single-copy components under pleiotropic constraint evolving by context-specific changes in regulation (undetectable with these methods but potentially elucidated by others), or 2) duplicating to escape pleiotropic constraint and ultimately producing sub-functionalized paralogs.

## Acknowledgments

Research reported here was supported by National Institute of General Medical Sciences of the National Institutes of Health under award number R35GM144001, from award #1939268 from the National Science Foundation Graduate Research Fellowship to OSZ, and by the National Science Foundation Postdoctoral Research Fellowships in Biology Program under award number DBI-2305817 to KDD.

